# Latent brain state dynamics predict early amyloid accumulation and cognitive impairment

**DOI:** 10.64898/2026.03.31.715655

**Authors:** Zhiyao Gao, Christina B. Young, Beyeongwook Lee, Rebecca E. Roush, Jay Kotulsky, Geraldine Cisneros, Elizabeth Mormino, Ann D. Cohen, Vinod Menon, Weidong Cai

**Affiliations:** Department of Psychiatry & Behavioral Sciences, Stanford University School of Medicine, Stanford, CA, USA; Department of Radiology, University of California Davis, Davis, CA, USA; Department of Neurology, University of California Davis, Davis, CA, USA; Department of Psychiatry, University of Pittsburgh, Pittsburgh, PA, USA; Department of Neurology & Neurological Sciences, Stanford University School of Medicine, Stanford, CA, USA; Wu Tsai Neuroscience Institute, Stanford University, Stanford, CA, USA; Maternal & Child Health Research Institute, Stanford, CA, USA

**Author notes:** Corresponding Authors : Zhiyao Gao; Vinod Menon; Weidong Cai.

**Keywords:** latent brain state, working memory, amyloid-β, mild cognitive impairment, fMRI

## Abstract

Amyloid-β (Aβ) accumulation is a continuous process central to pathological aging that begins decades before cognitive impairment emerges. While subthreshold Aβ levels have been linked to future decline in cognitive control, the neural mechanisms connecting this early accumulation to its neurocognitive impact are poorly understood. Brain circuit dynamics, which are essential for cognitive function, may offer a sensitive lens into these initial pathological changes. Here, we tested whether brain state dynamics could serve as sensitive markers for cognitive impairment at an early stage of Aβ burden. Using the Bayesian Switching Dynamic System (BSDS) model, we identified 4 distinct latent brain states from high-temporal-resolution (800 ms) fMRI data acquired from 116 older adults, including 72 cognitively normal (CN) individuals and 44 with mild cognitive impairment (MCI), during an N-back working-memory task. Adopting a dimensional approach, we examined how latent brain state dynamics relate to early amyloid burden, cognitive performance, and clinical symptoms. While Aβ levels failed to differentiate clinical groups or predict clinical symptoms and task performance, the dynamics of latent brain states proved highly sensitive to both early Aβ accumulation and cognition. Canonical correlation analysis revealed a significant relationship between brain state dynamics and early Aβ burden. Furthermore, the temporal properties of brain states were significantly predictive of working memory performance in CN individuals, a relationship that was selectively disrupted in the MCI group. The features of brain dynamics can also successfully predict cognitive impairment. Our findings establish brain state dynamics as sensitive neural markers of initial Aβ accumulation and early cognitive impairment, offering a new framework for developing predictive models to identify individuals at risk for future cognitive decline.

## Introduction

Dementia represents one of the most formidable public health challenges of our time, affecting more than 55 million people worldwide, a number projected to nearly triple by 2050 (Nichols et al., 2022; Organization, 2021). Despite decades of scientific progress, effective disease-modifying treatments remain elusive, largely because pathological changes accumulate decades before clinical symptoms emerge, by which point substantial and often irreversible neurodegeneration has already occurred (Becker & Greig, 2012; Becker et al., 2008; Cummings et al., 2020; Giacobini & Gold, 2013; Jack et al., 2013). A major goal for clinical trials and neuroscience is to understand the earliest neural alterations that precede overt cognitive decline, when the brain is still undergoing subtle pathological changes (Paraskevaidi et al., 2018; Sonnen et al., 2008). This early window encompasses a continuum from cognitively normal (CN) older adults to individuals with mild cognitive impairment (MCI), a transitional state that confers a substantially increased risk of progressing to dementia (Petersen et al., 2014). Characterizing this continuum provides a critical foundation for developing sensitive neural biomarkers that can identify at-risk individuals and guide timely intervention before irreversible damage occurs. Developing such biomarkers requires mechanistic insights into how early amyloid-β (Aβ) accumulation impacts neural circuit dynamics and, in turn, how these dynamics support cognitive function (Hampel et al., 2021; Hepp et al., 2016; Jansen et al., 2018; Takashima, 2009).

Aβ plaques is the first pathological feature of Alzheimer’s disease, and its accumulation represents a continuous process that begins decades before symptom onset (Hampel et al., 2021; Ingelsson et al., 2004; Sehar et al., 2022). Aβ preferentially deposits in cortical association hubs, such as the default mode network (DMN), and is associated with early disruptions in brain network connectivity (Bischof & Jacobs, 2019; Gallego-Rudolf et al., 2024; Hansson, 2021; Hyman et al., 2012; Palmqvist et al., 2017; Villeneuve et al., 2015). Aβ burden increases gradually, and even subthreshold levels below conventional PET-positivity cutoffs are now recognized as clinically meaningful (Elman et al., 2020; Ingelsson et al., 2004; Insel et al., 2016; Insel et al., 2017). In CN older adults, subtle Aβ burden predicts future decline in executive function more accurately than baseline clinical status (Ali et al., 2022; Bischof & Jacobs, 2019; Chen et al., 2025; Tideman et al., 2022). Critically, a recent study reported that the rate of Aβ accumulation in frontoparietal regions, key nodes of the brain’s cognitive control network, specifically predicted longitudinal decline in executive function (Chen et al., 2025). This aligns with growing evidence that disruptions in cognitive control, encompassing attention, working memory, and inhibitory regulation, emerge early in the disease process (Amieva et al., 2004; Baddeley et al., 2001; Germano & Kinsella, 2005; Kaiser et al., 2018; Li et al., 2015; Martyr et al., 2019; Storandt, 2008). Such impairments can significantly affect everyday functioning and are among the most reliable behavioral predictors of future cognitive decline (Clément et al., 2013; Saunders & Summers, 2011; Storandt et al., 2006; Summers & Saunders, 2012). However, the neural mechanisms through which early Aβ accumulation disrupts cognitive control processes remain poorly understood.

Cognitive control refers to the fundamental ability to coordinate cognitive and motor processes in accordance with current and future goals (Hammond & Summers, 1972; Menon & D’Esposito, 2022; Miller, 2000). Cognitive control relies on the dynamic interaction of distributed networks (Cocchi et al., 2013; Cole & Schneider, 2007; Menon & D’Esposito, 2022; Power & Petersen, 2013), including the frontoparietal network (FPN) (Fassbender et al., 2006; Gao et al., 2021; Nee, 2021; Zanto & Gazzaley, 2013); the salience network (SN) (Chen et al., 2016; Seeley, 2019; Uddin, 2015); and the default mode network (DMN) (Anderson et al., 2016; Menon, 2023; Smallwood et al., 2021; Wang et al., 2021). Brain states and their temporal evolution are crucial concepts for understanding how rapid changing and coordinated neural activity supports cognitive function (Greene et al., 2023; Taghia et al., 2018). One prominent hypothesis is that timely engagement and maintenance of task-optimal brain states facilitate the allocation of attention and cognitive resource to goal-directed processes (Cai et al., 2024; Cai et al., 2021; Lee et al., 2022; Taghia et al., 2018). Disruptions to these dynamics, such as a poor ability to disengage from non-optimal brain states, are increasingly recognized as a key feature of cognitive impairment in both aging and neuropsychiatric conditions (Cai et al., 2023; Cai et al., 2024; Cai et al., 2021; Lee et al., 2022; Taghia et al., 2018). However, it remains poorly understood how the brain state dynamics are impacted by the initial stages of Aβ accumulation.

To address these challenges, we analyzed data from 116 participants with MCI and matched CN controls from the HCP Connectomics in Brain Aging and Dementia (HCP-CBA) study (Cohen et al., 2021). The dataset included clinical, cognitive, PET, and task-fMRI (n-back working memory) data. PET was used to estimate Aβ burden. Notably, most of the participants had subthreshold Aβ levels. Our primary aim was to determine how Aβ burden impacts cognitive function, brain activation, and brain state dynamics. We also sought to identify differences in these measures between the MCI and CN groups. To investigate brain dynamics, we utilized Bayesian Switching Dynamic System (BSDS), a state-space model that identifies latent brain states and their temporal evolution based on unique activation and connectivity patterns (**Figure 2C**) (Taghia et al., 2018). This method overcomes the limitations of conventional sliding window approaches, such as rigid temporal boundary and arbitrarily model parameters (Hutchison et al., 2013; Taghia et al., 2018). This rigorous computational framework allowed us to link brain state dynamics to early Aβ burden, cognitive function and clinical symptoms. We hypothesized that even subthreshold Aβ burden would significantly impact brain state dynamics and that these dynamics would, in turn, predict cognitive performance and clinical symptoms.

## Results

### Participants

We analyzed data from the HCP Connectomics in Brain Aging and Dementia (HCP-CBA) study (Cohen et al., 2021), which included 72 CN participants (48 female, 24 male; 63.83 years; 8 Aβ+, 64 Aβ-) and 44 individuals with MCI (32 female, 12 male; mean age 62.21 years; 4 Aβ+, 40 Aβ-), all of whom completed two runs of the N-back task (**Table 1**). The CN and MCI groups were matched for age, gender, and head motion (all ps > 0.32 for the full sample and all ps > 0.28 for the fMRI + behavior sample; see **Table 1-2**).

**Table 1.** Demographics (fMRI sample)

| **Demographics** | **CN (N=72)** | **MCI (N = 44)** | **p-value** |
| --- | --- | --- | --- |
| Age (SD) | 63.833(8.222) | 62.214(8.595) | 0.321 |
| Gender (Female/Male) | 48/24 | 32/12 | 0.494 |
| MOCA (SD) | 25.972(4.049) | 23.727(2.582) | 0.001 |
| Head Motion (SD) | 0.174(0.056) | 0.165(0.050) | 0.405 |
| SUVRs (SD) | 1.193(0.259) | 1.155(0.219) | 0.452 |

CN: cognitively normal; MCI: mild cognitive impairment

As expected, compared to the CN group, the MCI group showed significantly lower Montreal Cognitive Assessment (MoCA) scores, indicating measurable cognitive impairment (full sample: p = 0.001; fMRI + behavior sample: p = 0.003; **Table 1-2**). Notably, the low prevalence of Aβ+ in both the CN (11.1%) and MCI (9.1%) groups indicates that Aβ pathology is not the primary differentiator between these clinical groups.

Behavioral data from 14 participants were missing, resulting in a final sample of 102 participants for the fMRI and behavioral analysis. The fMRI + behavioral sample included 66 CN participants (44 female, 22 male; 63.47 years; 7 Aβ+), 36 MCI patients (27 female, 9 male; 62.57 years; 1 Aβ+) (**Table 2**).

**Table 2.** Demographics (fMRI + behavior sample)

| **Demographics** | **CN (N=66)** | **MCI (N = 36)** | **p-value** |
| --- | --- | --- | --- |
| Age (SD) | 63.470(7.671) | 62.571(8.846) | 0.597 |
| Gender (Female/Male) | 44/22 | 27/9 | 0.382 |
| MOCA (SD) | 26.046(4.178) | 23.722(2.722) | 0.003 |
| Head Motion (SD) | 0.177(0.057) | 0.165(0.051) | 0.281 |
| SUVRs (SD) | 1.172(0.225) | 1.175(0.257) | 0.950 |

CN: cognitively normal; MCI: mild cognitive impairment

### MCI participants show substantial working memory deficits

We conducted behavioral analyses to investigate the impact of working memory (WM) load on task performance and to assess group differences between CN and individuals with MCI. WM accuracy was analyzed using a 2 (Group: CN, MCI) × 2 (WM Load: 0-back, 2-back) mixed-design ANOVA. While there was no significant Group × Load interaction (F(1,100) = 0.189, p = 0.664, η^2^_p_ = 0.002), a significant main effect of WM Load emerged: participants were more accurate on the 0-back task than on the 2-back task (F(1,100) = 9.234, p = 0.003, η^2^_p_ = 0.085). Moreover, CN exhibited higher overall accuracy compared to MCI (main effect of Group: (F(1,100) = 12.476, p < 0.001, η^2^_p_= 0.111) (**Figure 1**).

**Figure 1.**
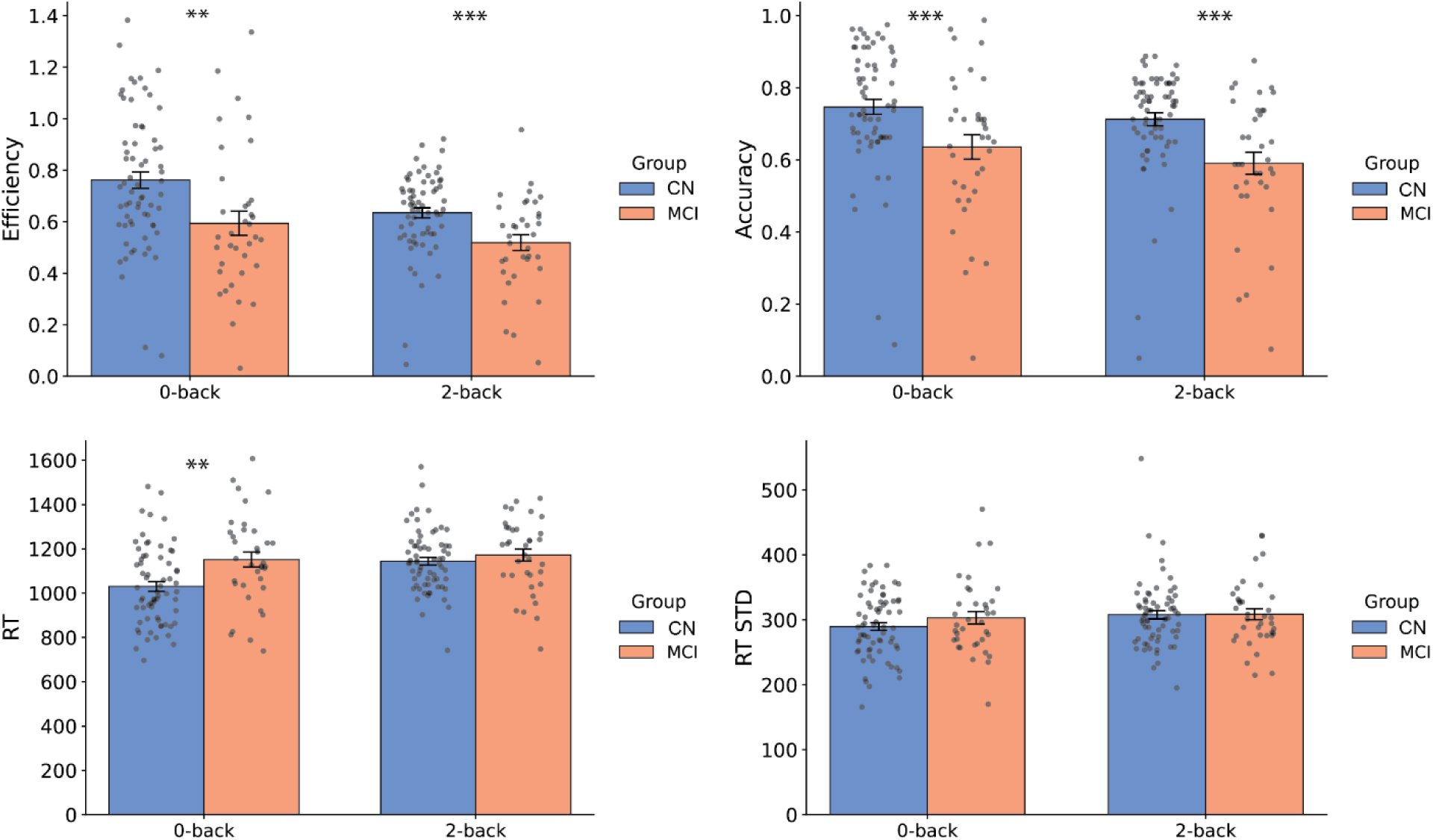
Reduced behavioral efficiency, accuracy, mean reactive time, and response stability (standard deviation of reaction time) in individuals with MCI. * indicates p < 0.05, *** indicates p < 0.001.

A similar 2 x 2 ANOVA on mean reaction time (RT) showed significant effect of interaction (F(1, 100) = 11.129, p = 0.001, η^2^_p_ = 0.100) and a main effect of WM load (F(1,100) = 23.292, p < 0.001, η^2^_p_ = 0.189), as well as a significant main effect of group (F(1,100) = 5.352, p = 0.023, η^2^_p_ = 0.051). Simple effect analysis revealed that CN responded faster than MCI in 0-back trials (p = 0.002) but did not show significant difference in 2-back trials (p = 0.258).

Behavioral efficiency measures how well participants can achieve high accuracy while minimizing the reaction time required to do complete the task, providing a more holistic understanding of task performance. Two-way ANOVA revealed higher efficiency in CN than MCI (F(1,100) = 11.587, p < 0.001, η^2^_p_ = 0.104), and in 0-back trials compared to 2-back trials (F(1,100) = 24.969, p < 0.001, η^2^_p_ = 0.200), with no significant interaction (F(1,100) = 1.646, p < 0.203, η^2^_p_ = 0.016).

Intra-individual response variability (IIRV) was assessed using the standard deviation (STD) of reaction times for each condition. A two-way ANOVA on STD revealed no significant interaction effect between group and load (F(1, 100) = 0.858, p = 0.356, η^2^_p_= 0.009), nor a main effect of group (F(1,100) = 0.734, p = 0.393, η^2^_p_ = 0.007). There was only a marginal main effect of Load (F(1, 100) = 3.120, p = 0.080, η^2^_p_= 0.030). Simple effects analysis did not find significant differences of response variability either on STD for both the 0-back and the 2-back condition (ps > 0.705) (**Table 3**), indicating comparable level of response variability across groups.

**Table 3.**
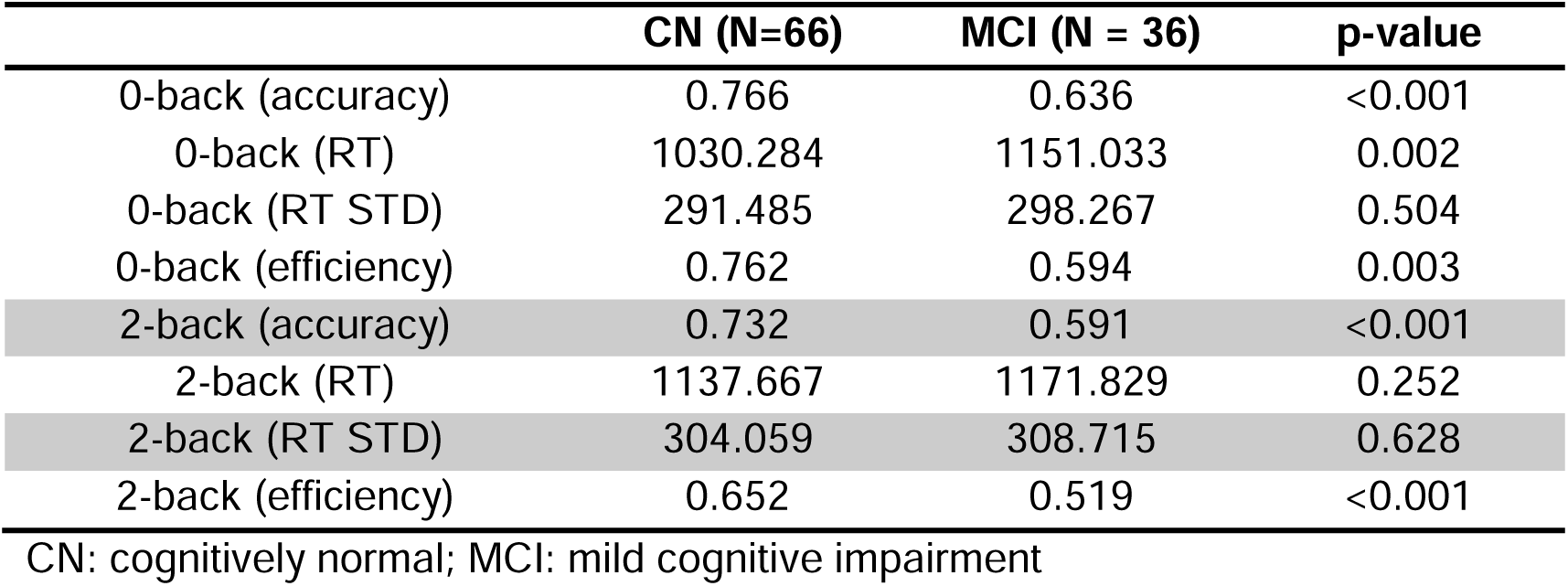
Behavioral performance (fMRI + behavior sample)

|  | **CN (N=66)** | **MCI (N = 36)** | **p-value** |
| --- | --- | --- | --- |
| 0-back (accuracy) | 0.766 | 0.636 | <0.001 |
| 0-back (RT) | 1030.284 | 1151.033 | 0.002 |
| 0-back (RT STD) | 291.485 | 298.267 | 0.504 |
| 0-back (efficiency) | 0.762 | 0.594 | 0.003 |
| 2-back (accuracy) | 0.732 | 0.591 | <0.001 |
| 2-back (RT) | 1137.667 | 1171.829 | 0.252 |
| 2-back (RT STD) | 304.059 | 308.715 | 0.628 |
| 2-back (efficiency) | 0.652 | 0.519 | <0.001 |

CN: cognitively normal; MCI: mild cognitive impairment

In summary, CN demonstrated consistently higher accuracy, faster responses, and higher efficiency than MCI, with WM load modulated task performance for both CN and MCI.

### Amyloid-β accumulation fails to predict cognitive performance

Having established clear behavioral differences between groups, we next examined whether conventional biomarkers could account for these cognitive deficits. Standardized uptake value ratios (SUVRs) across the brain did not significantly differ between CN and MCI (full sample: p = 0.45; fMRI + behavior sample: p = 0.95; **Table 1-2**; and see **Figure 2** for regional SUVRs).

**Figure 2.**
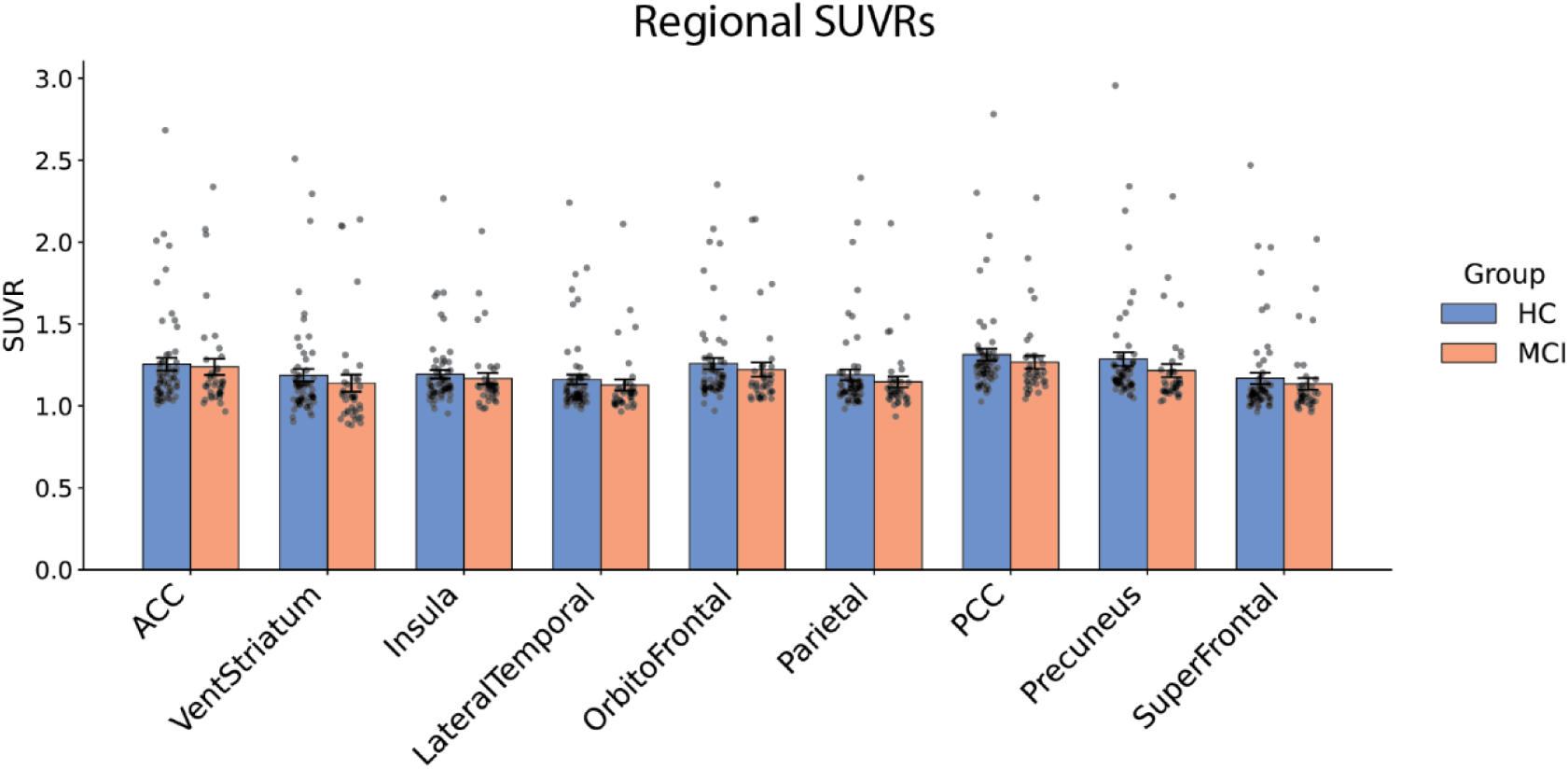
Regional SUVRs in the brain. ACC, anterior cingulate cortex; VentStriatum, ventral striatum; LateralTemporal, lateral temporal lobe; Orbitofrontal, orbital frontal cortex; PCC, posterior cingulate cortex; SuperFrontal, superior frontal gyrus.

Using a dimensional approach that pools CN and MCI participants, we then assessed the relationship between amyloid burden and clinical symptoms as indexed by MoCA scores, which revealed no significant associations after controlling for age and gender (all p > 0.647, FDR corrected across 10 regions of interest: 9 regional and 1 global SUVRs). We then examined whether amyloid burden was associated with cognitive performance on the 0-back and 2-back tasks adopting a dimensional approach. No significant associations were found between amyloid burden and behavioral measures—including accuracy, reaction time (RT), RT variability (RT SD), or efficiency after controlling for age and gender, whether using regional or global SUVRs (all p > 0.098, FDR-corrected across 10 SUVRs). Collectively, these findings provide converging evidence that our participants are in an early phase of amyloid-β burden that do not differentiate between CN and MCI participants.

We further explored potential multivariate associations between regional Aβ burden and combined measures of task performance and clinical symptoms using canonical correlation analysis (CCA) across all participants, while controlling for age and sex. The analyses included (1) task performance metrics including accuracy, RT, and RT standard deviation (STD) from both 0-back and 2-back conditions along with MoCA scores; (2) task performance from both 0-back and 2-back conditions only; and (3) task performance from either the 0-back or 2-back condition individually. However, no significant canonical correlations were observed in any of these analyses (all ps > 0.989, FDR-corrected).

These findings demonstrate that in this non-demented cohort with predominantly subthreshold amyloid levels, Aβ burden failed to differentiate clinical groups, predict cognitive symptoms, or account for behavioral performance.

### Amyloid-β burden does not predict brain activity levels

We investigated the relationship between regional/global Aβ burden (SUVRs) and task-related brain activation, controlling for age, sex, and head motion. Task activation was quantified using a traditional univariate GLM, where 0-back and 2-back conditions were modeled as separate regressors. The resulting parameter estimates for each condition were then extracted from 19 ROIs derived from the BSDS analysis for our primary categorical and secondary dimensional analysis.

For the categorical analysis, a repeated-measures ANOVA was conducted with group as a between-subject factor and working memory (WM) load and ROI as within-subject factors. No significant categorical differences were observed. Specifically, there was no main effect of group (F(1,100) = 1.485, p = 0.226, η^2^_p_ = 0.015), nor were there significant interactions involving group (group by WM by ROI: F(7.165, 1126.119) = 0.636, p = 0.752, η^2^_p_ = 0.006; group by WM: F(1, 100) = 0.083, p = 0.774, η^2^_p_ = 0.001; group by ROIs: F(10.499, 1049.865) = 0.600, p = 0.823, η^2^_p_ = 0.006), indicating no categorical differences between CN and MCI groups. In contrast, significant effect were observed for interaction between WM and ROIs (F(8.202, 1126.119) = 11.919, p < 0.001, η^2^_p_ = 0.106), and main effects of WM (F(1, 100) = 22.875, p < 0.001, η^2^_p_ = 0.186) and ROIs (F(10.499, 4096.860) = 68.845, p < 0.001, η^2^_p_ = 0.408), reflecting robust WM load effects and region-specific activation differences across participants.

In the dimensional analysis, no significant associations were found between either regional or global SUVRs and task-related activation in either condition (0-back: all ps > 0.126, FDR-corrected; 2-back: all ps > 0.358, FDR-corrected). To further assess potential multivariate relationships, we conducted a canonical correlation analysis (CCA) between nine regional SUVRs and activation across the 19 ROIs during task performance, adjusting for age, sex, and head motion. This analysis revealed no significant associations between activation patterns and regional SUVRs (0-back: p = 0.114; 2-back: p = 0.143; FWE-corrected).

These results indicate that early Aβ burden was not significantly associated with static task-related activation in this cohort, and underscore the need for more sensitive functional markers that can capture the neural basis of emerging cognitive deficits.

### Brain state dynamics during working memory

To understand the dynamic neural processes underlying working memory performance, we examined latent brain state dynamics across 19 regions of interest (ROIs) from four key networks: the frontoparietal network (FPN), salience network (SN), and default mode network (DMN) (Menon & D’Esposito, 2022). They encompass bilateral anterior insula (AI) and pre-supplementary motor area (pre-SMA or dorsomedial prefrontal cortex, DMPFC) from the SN, bilateral middle frontal gyrus (MFG), frontal eye fields (FEF), and inferior parietal lobe (IPL) from the FPN, posterior cingulate cortex (PCC) and ventromedial prefrontal cortex (VMPFC) from the DMN, and 4 category-specific visual ROIs from each hemisphere, each selective for one of the four categories employed in the n-back task (i.e. face, place, body, and tools) (**Figure 3A**) (see Methods).

**Figure 3.**
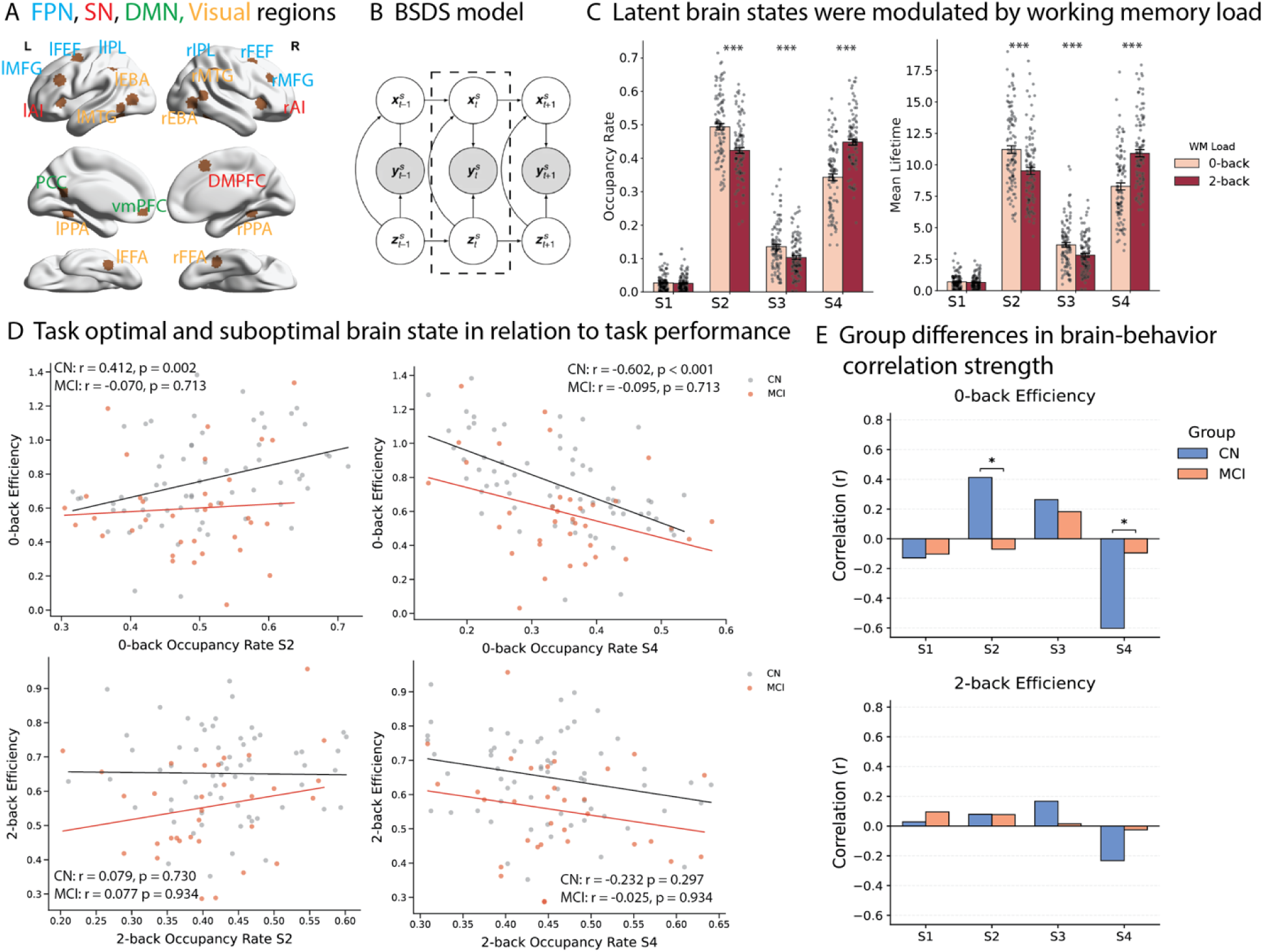
Latent brain state dynamics (mean lifetime) during WM and their relationship to task performance. A. Key regions implicated in cognitive control: frontal-parietal network: bilateral middle frontal gyrus (MFG), frontal eye field (FEF), inferior parietal lobe (IPL); salience network (SN): bilateral anterior insular (AI), dorsal medial frontal cortex (dmPFC); and default mode network (DMN): ventral medial frontal cortex (vmPFC) and posterior cingulate cortex (PCC), as well as visual selective areas in both hemispheres: face fusiform area (FFA), place parahippocampus area (PPA), extrastriate body area (EBA), and tool selective area in middle temporal gyrus (MTG). B. BSDS model: Bayesian switching dynamics states model. C. Latent brain state dynamics were modulated by working memory load. D. Correlation between latent brain state dynamics and task performance in CN and MCI after accounting for age, gender, and head motion. Under 0-back condition, occupancy rate of state S2 was positively associated with efficiency; S4 was negatively associated with efficiency in cognitively normal participants but not MCI. Under 2-back condition, occupancy rate of S2 and S4 did not show significant association with behavior efficiency in both groups. E. Group differences in brain state–behavior coupling in 0-back and 2-back tasks. S2 and S4 occupancy rates show stronger coupling with efficiency in CN than MCI during the 0-back task. * indicates p < 0.05, *** indicates p < 0.001.

We applied the Bayesian Switching Dynamic System (BSDS) model (Taghia et al., 2018) to identify latent brain states from high-temporal-resolution (800 ms) fMRI data. BSDS is a generative state-space model that identifies discrete brain states, each characterized by unique patterns of activation and connectivity, and tracks how the brain transitions between these states over time (**Figure 3B**). Unlike conventional sliding window approaches that impose arbitrary temporal boundaries, BSDS uses a principled statistical framework to detect state transitions based on the data itself. Applying this model to the combined cohort of CN and MCI participants, we identified 4 distinct brain states, labeled S1 through S4 (**Figure 3C**).

For each brain state, we quantified two key temporal properties that capture how the brain engages and maintains functional configurations. Occupancy rate measures how often a latent brain state occurs, reflecting the proportion of time the brain spends in that particular state. Mean lifetime measures the average duration for which a brain state persists before transitioning to another state, offering insights into the stability of that functional configuration. Together, these metrics characterize the temporal dynamics of brain state engagement during cognitive performance. Additional analyses revealed that S2 has highest occupancy rate and mean lifetime in 0-back, while S4 has highest occupancy rate and mean lifetime in 2-back (**Figures 3C**) and demonstrate that specific brain states are differentially sensitive to working memory demands (see SI results).

*MCI vs. CN differences in latent state dynamics* To determine if MCI differed from CN in dynamic temporal state properties, we conducted separate two-way mixed ANOVAs for mean lifetime and occupancy rate with Group (MCI, CN) as a between-subjects factor, and Latent Brain State (S1-S4) as within-subjects factor. ANOVA for mean lifetime revealed a significant main effect of State (F(2.52, 287.66) = 855.049, p<0.001, η^2^_p_ = 0.882, Greenhouse-Geisser corrected). However, we found no significant interaction between Group and State (F(2.52, 287.66) = 1.000, p = 0.393, η^2^_p_ = 0.009, Greenhouse-Geisser corrected), and no main effect of group (F(1,114) = 0.072, p = 0.788, η^2^_p_ = 0.001). Similarly, ANOVA for occupancy rate (S2-S4, S1 was excluded to avoid collinearity issue as the sum of 4 occupancy rate will be 1) showed a significant main effect of brain state (F(1.797, 204.887) = 271.151, p<0.001, η^2^_p_ = 0.704, Greenhouse-Geisser corrected), but there was no significant interaction between Group and State (F(1.797, 204.887) = 1.61, p = 0.205, η^2^_p_ = 0.014, Greenhouse-Geisser corrected), or main effect of group (F(1,114) = 0.072, p = 0.789, η^2^_p_ = 0.001). These results suggest that, in pre-dementia stages, MCI and CN participants exhibit comparable group-level temporal properties of latent brain states.

### Brain state dynamics predict task performance in CN but not in MCI

The finding that MCI and CN participants generate similar brain states that are appropriately modulated by task demands raises a critical question: if both groups access the same functional configurations, why do MCI participants show substantial behavioral deficits? To address this, we examined whether MCI participants effectively utilize these states for successful task performance.

In CN, during 0-back, longer lifetimes of S2 and S3 correlated positively with efficiency (r = 0.393, 0.275; p = 0.003, 0.041) and accuracy (r = 0.459, 0.293; p < 0.001, 0.031), and with faster responses (r = −0.260, −0.324; p = 0.053, 0.020). Longer S4 lifetimes were associated with lower efficiency (r = −0.582; p < 0.001) and accuracy (r = −0.579; p < 0.001), slower responses (r = 0.503; p < 0.001), and greater variability (r = 0.357; p = 0.016). Occupancy rates mirrored these effects: greater S2 occupancy correlated with higher efficiency (r = 0.412; p = 0.002) and accuracy (r = 0.462; p < 0.001); greater S3 occupancy correlated with faster responses (r = −0.331; p = 0.017). In contrast, greater S4 occupancy predicted lower efficiency (r = −0.602; p < 0.001) and accuracy (r = −0.624; p < 0.001), and slower responses (r = 0.489; p < 0.001). In the 2-back condition, only S4 dynamics were significantly related to slower responses (lifetime: r = 0.373; p = 0.012; occupancy: r = 0.354; p = 0.022) (**Figure 3D**). Strikingly, in MCI, no brain–behavior associations survived FDR correction under either load condition (all p > 0.342) (**Figure 3D**).

To assess whether associations between brain state dynamics and behavioral performance differed by group, we compared partial correlation coefficients between CN and MCI participants using Fisher’s r-to-z transformations, chosen to account for potential confounds such as age, sex, and head motion that may influence both brain measures and behavior. For mean lifetime, State S4 showed a more negative association with efficiency (Z = −2.88, p = 0.016) in CN relative to MCI. For occupancy rate, the analysis revealed a significantly stronger positive association between S2 and efficiency in the CN group compared with the MCI group (r: 0.412 vs -0.070; Z = 2.37, p = 0.036), as well as a significantly stronger negative association between S4 and efficiency in CN than in MCI (r: -0.602 vs.-0.065; Z = −2.80, p = 0.021) during 0-back trials. All reported p values were corrected for multiple comparisons using false discovery rate (FDR) correction (**Figure 3E**).

These results suggest that cognitively normal individuals show that when the non-optimal brain state (S4, optimized for 2-back) is engaged during the 0-back task, performance suffers. This adaptive brain-behavior coupling is absent in MCI, pointing to a failure to effectively utilize brain states to support working memory performance.

### Amyloid-β burden impairs brain state dynamics

We examined whether individual differences in early amyloid-β burden across brain regions impairs brain state dynamics, even in this predominantly subthreshold cohort where Aβ burden failed to predict cognition.

We used CCA to test whether the temporal features of brain states (mean lifetime and occupancy rate) could jointly predict Aβ burden across nine brain regions in the entire sample, controlling for age, gender, and head motion. This multivariate analysis allowed us to examine the complex relationship between the temporal features of brain states (i.e. mean lifetime and occupancy rate) and multi-regional Aβ levels. CCA revealed a significant relationship between the temporal features of latent brain states even after covarying for age, gender, and head motion (r=0.580, p=0.007; FWE corrected). (**Figure 4A**).

**Figure 4.**
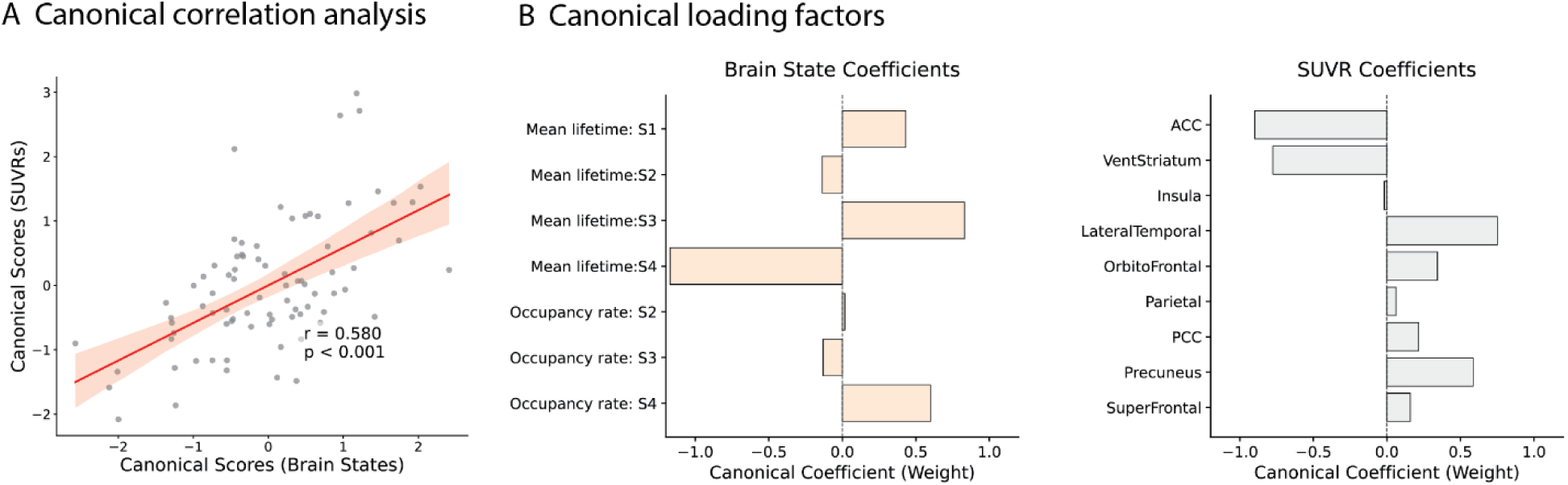
Latent brain state dynamics predict Aβ accumulation. A. Canonical correlation between latent brain state dynamics and regional SUVRs. B. Canonical loading factors. ACC, anterior cingulate cortex; VentStriatum, ventral striatum; LateralTemporal, lateral temporal lobe; Orbitofrontal, orbital frontal cortex; PCC, posterior cingulate cortex; SuperFrontal, superior frontal gyrus.

The CCA model was primarily driven by opposing contributions from brain state dynamics. Positive CCA coefficients were observed for S3 lifetime and S4 occupancy rate, while negative coefficients were found for state S2 (both lifetime and occupancy rate) and S4 mean lifetime. This model was associated with widespread positive CCA coefficients across most SUVR regions, with the exception of the anterior cingulate cortex (ACC), ventral striatum, and insula (**Figure 4B**).

Canonical loading analysis revealed the strength of association between individual variables and the canonical variates. Significant associations were found for the mean lifetimes of S1 (r = 0.263, p = 0.030), S3 (r = 0.269, p = 0.030), S2 (r = –0.309, p = 0.016), S4 (r = –0.503, p < 0.001). Significant canonical loadings were also observed in multiple cortical regions, including the insula (r = 0.308, p = 0.005), lateral temporal lobe (r = 0.452, p < 0.001), orbitofrontal cortex (r = 0.353, p = 0.001), parietal cortex (r = 0.434, p < 0.001), posterior cingulate cortex (r = 0.377, p < 0.001), precuneus (r = 0.498, p < 0.001), and superior frontal gyrus (r = 0.456, p < 0.001). All p-values were corrected for multiple comparisons using FDR, and Pearson’s correlation was used for all analyses.

These results demonstrate that brain state dynamics are sensitive to early amyloid accumulation even when Aβ levels are predominantly subthreshold and fail to predict cognitive outcomes.

### Brain state dynamics predict clinical impairment

Having demonstrated that brain state dynamics predict task performance in CN individuals and that this relationship is disrupted in MCI, we next examined the associations between brain-state dynamics, Aβ accumulation, and cognitive impairments (MoCA scores). We trained a series of Lasso regression models to predict MoCA scores, with performance evaluated via leave one out cross validation (LOOCV).

A model using only Aβ levels from regional SUVRs as features did not significantly predict MoCA scores (r = -0.067, p = 0.514). In contrast, a model based on brain dynamic features (occupancy rates of brain states during the 0-back condition) yielded significant predictions (r = 0.330, p = 0.003) (**Figure 5**). An analysis of this model’s features revealed that the top predictors for MoCA scores were occupancy rates of S2 and S3, whereas S4 occupancy exhibited a weak negative influence (see Supplementary Figure S3).

**Figure 5.**
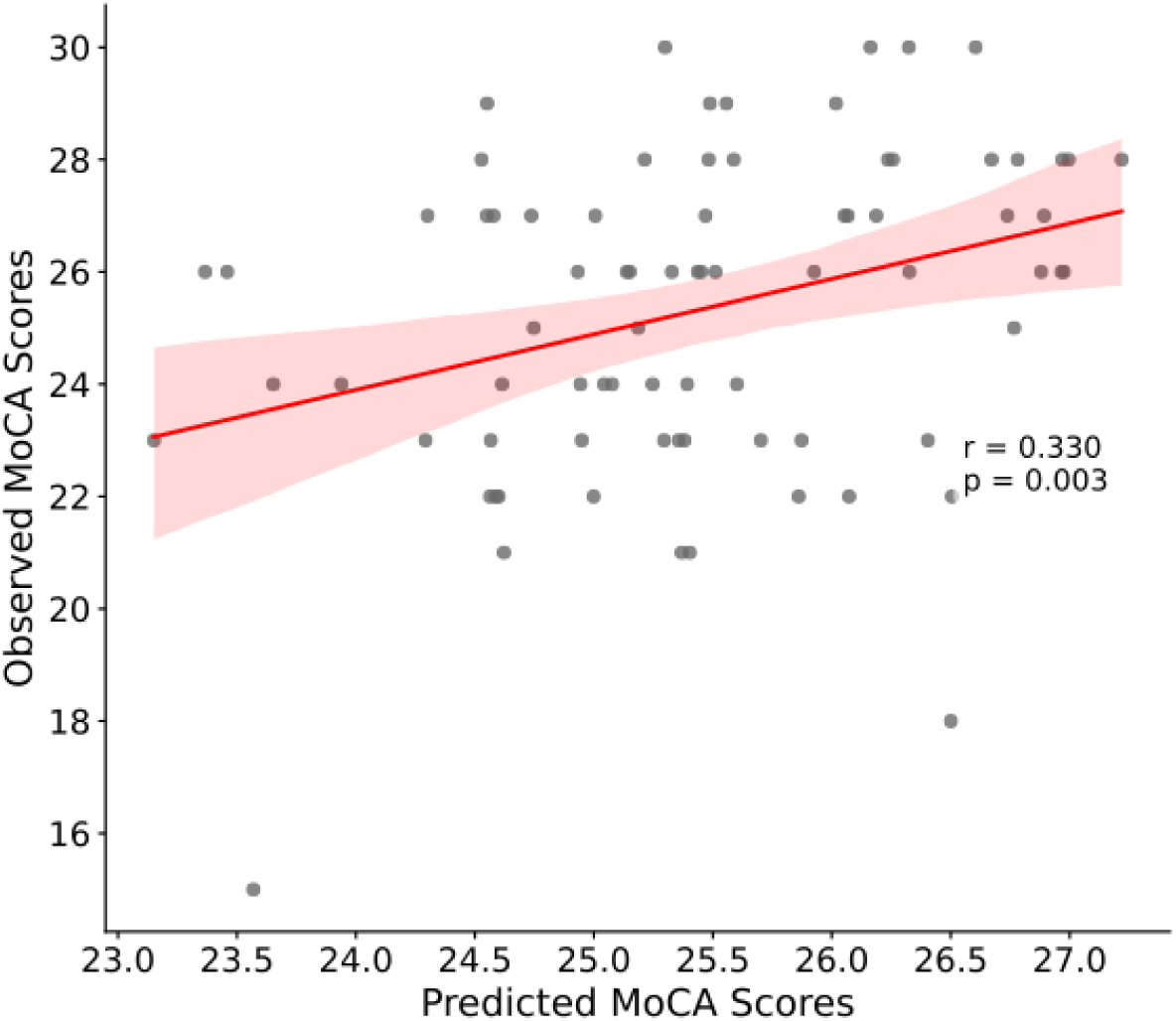
Predicting MoCA Scores with Brain Dynamics. A Lasso regression model using brain state occupancy rates from the 0-back condition as features significantly predicted MoCA scores (p = 0.003).

Together, these findings suggest that latent brain state dynamics may serve as a sensitive biomarker of cognitive impairment in elders with sub-threshold Aβ levels where Aβ burden alone is not yet predictive.

## Discussion

One of the major challenges in neurodegeneration research is understanding the neural mechanisms connecting early, subthreshold Aβ accumulation to its neurocognitive impact. Here we tested the hypothesis that brain state dynamics could serve as a sensitive neural marker for identifying this early neural impact. Applying the Bayesian switching dynamical system (BSDS) model to high-temporal resolution (TR = 800 ms) fMRI data, we identified 4 latent brain states in the cohort of older adults with normal cognition and MCI when they were engaged in the n-back working memory task. While early Aβ accumulation failed to predict clinical symptom, task performance and brain activation and even to differentiate clinical groups, brain state dynamics are sensitive to individual differences in early Aβ burden. Consistent with previous findings of young adults, brain state dynamics were also associated with behavioral performance in old adults. Critically, these dynamic features are predictive of working memory performance in CN individuals, a relationship that was absent in the MCI group. Finally, prediction models using brain dynamic features successfully predict individuals’ clinical assessment score of cognitive impairment, establishing them as a robust functional biomarker.

### Amyloid level as a poor predictor in the subthreshold range

Previous studies have shown that, even below the PET-positivity threshold, Aβ levels can indicate individuals at risk for future accumulation, cognitive decline, and emergent tau pathology (Bischof & Jacobs, 2019). This presents an opportunity to develop clinically useful biomarker for identify elders that may require preventative intervention before any clinical symptoms can be observed. However, the extent to which subthreshold Aβ levels impacts neurocognitive function remain unknown. Our study demonstrated that, in an older adult cohort with predominantly subthreshold Aβ levels, amyloid burden itself is a poor predictor of cognitive and clinical status. Specifically, the multi-regional SUVRs failed to predict behavioral performance, regional activation in the n-back task, or MoCA scores. This highlights a critical gap: conventional behavioral and neural markers may struggle in this early, preclinical phase where Aβ accumulation is at low level or this cognitive impairment in this cohort may not be driven by amyloid pathology.

Moreover, we did not find significant differences in Aβ levels between the CN and MCI groups. This absence of a group difference may reflect heterogeneous neurodegenerative processes in the MCI group. With only 4 MCI participants meeting the PET threshold of amyloid positivity (Cohen et al., 2021; Cohen et al., 2013), it is likely that a substantial proportion of the MCI group may have non-AD etiologies, such as vascular cognitive impairment , Lewy body disease, or other mixed pathologies (Harding & Halliday, 2001; Jellinger & Attems, 2010; Ossenkoppele et al., 2015). This heterogeneity underscores the critical need for the dimensional approach we adopted, which can capture continuous relationships that categorical comparisons would miss.

### Brain state dynamics predict early amyloid level despite its weak behavioral links

A major finding of this study is that brain state dynamics, unlike trial-averaged activation, proved sensitive to early amyloid accumulation. Our canonical correlation analysis (CCA) revealed a significant multivariate association between the temporal properties of brain states (i.e. mean lifetime, occupancy rate) and regional Aβ deposition. This relationship remained significant after accounting for demographic and motion covariates, underscoring its robustness. In sharp contrast, similar analyses failed to identify any significant multivariate association between trial-averaged brain activation and regional Aβ deposition. This suggests that, while subthreshold amyloid levels do not yet have an observable effect on overall task-induced activation, this early Aβ accumulation already impacts more subtle brain state dynamics. Further canonical loading analyses highlighted that the load-dominated states (S2 and S4) and early amyloid accumulation contributed in opposing directions to this association. This indicates that early amyloid accumulation may disrupt the brain’s efficient transition to or maintenance of the brain state that is optimal to meet a given cognitive demand.

### Load dependent brain dynamics in older adults

State-space models have been increasingly used to understand brain dynamics and their association with cognition (Greene et al., 2023; Taghia et al., 2018). Unlike traditional activation or connectivity measures that provide static summaries of brain activity, state-space models capture how the brain dynamically transitions between functional configurations over time, a process that is essential for the flexible deployment of cognitive control (Cai et al., 2024; Cai et al., 2021; Lee et al., 2022; Menon & D’Esposito, 2022; Taghia et al., 2018). Using the BSDS, our previous study revealed distinct brain states when a cohort of young adults was engaged in the n-back working memory task; importantly, individuals’ ability to engage and maintain a task-optimal brain state was tightly associated with their working memory performance (Taghia et al., 2018). Here, we replicated key aspects of these findings in a cohort of older adults. BSDS identified four latent brain states that were differentially engaged by task demands: S2 was dominant during the 0-back condition, while S4 was more prominent during the 2-back condition. This demonstrates a dynamic recruitment of brain states in response to changing cognitive control demands. We also found that the occupancy rate of state S2 predicted behavioral measures of the 0-back task, such as accuracy and RT, replicating our previous findings (Taghia et al., 2018). This implies that S2 is an optimal state for the 0-back task, where its engagement facilitates task performance.

A key finding, however, was a critical divergence from our work in younger adults regarding the high-load 2-back condition. While S4 was clearly the dominant state during the 2-back task, its dynamic features lacked a strong association with behavioral performance. This is in direct contrast to our previous study, where the dominant 2-back state was strongly predictive of performance. This discrepancy does not appear to be because S4 is the “wrong” state. Its neural profile, greater activation in frontoparietal and salience networks and greater DMN deactivation compared to S2, is precisely what is expected of a high-load, cognitively demanding state (Supplementary Results and Figure S1-2). This pattern is consistent with our findings in young adults (Taghia et al., 2018) and aligns with extensive evidence showing that increasing working memory load enhances activation in frontoparietal and salience regions while suppressing DMN activity (Gao et al., 2021; Kirschen et al., 2005; Manoach et al., 1997; Owen et al., 2005; Wang et al., 2019). Instead, our findings point to an age-related inefficiency in the use of this state. First, the overall occupancy rates and mean lifetimes of all brain states were lower in this older cohort relative to the previous findings in the young cohort (Taghia et al., 2018), suggesting a general age-related effect on brain state stability. Second, the engagement of this high-load state (S4) during the simple 0-back task was associated with poorer performance. This suggests that recruiting S4 is inefficient and potentially maladaptive during low-demand conditions. Therefore, while older adults can recruit the correct high-load state, they may be unable to engage or maintain it as effectively as younger adults.

### Brain state dynamics predict behavioral performance in CN but not in MCI

Another important finding of this study is the profound dissociation in brain-behavior relationships between the CN and MCI groups. In CN participants, brain state dynamics were linked to behavioral performance in the n-back working memory task. For example, greater occupancy rate and longer mean lifetimes of the low-load dominated state (S2) were strongly associated with higher accuracy and faster response times on the 0-back trials. Conversely, engagement of the high-load dominated state (S4) during the same low-load condition predicted poorer performance. This pattern suggests that the S2 represents an optimal state for performing the 0-back task, while engaging the “high-load” S4 state indicates that these individuals were struggling or processing the task inefficiently.

Strikingly, this adaptive relationship was absent in the MCI group. Despite generating a similar dynamic patterns of latent states, their temporal properties showed no significant association with behavioral performance. This null finding provides a critical insight. It suggests that a core deficit in MCI may not be the inability to access certain neural states, but rather the failure to utilize these states effectively. While the MCI brain may still “visit” the correct states, this engagement is inefficient and no longer translates into successful cognitive action. This lack of association itself may serve as a highly sensitive marker of emerging cognitive impairment, reflecting a breakdown in the brain’s ability to tune its dynamics to optimize behavior. This interpretation is consistent with evidence that large-scale connectivity patterns remain relatively preserved in aging (Adams et al., 2023; Cabeza et al., 2018; Stern et al., 2020) and resonates with the concept of aging involving a decline in cognitive flexibility and an emerging inefficiency in neural processing (Lee et al., 2022), reflecting a diminished capacity of the brain to dynamically orchestrate its states to meet task demands.

### Brain state dynamics as powerful predictor of cognitive impairment

Our findings established brain state dynamics as a powerful and sensitive predictor of cognitive impairments in a cohort of elder adults whose amyloid levels are mostly below a positivity threshold. We trained prediction models using brain dynamics as features that predict a widely used screening test for cognitive impairment, i.e. MoCA. A Lasso regression model trained on state occupancy rates during the 0-back condition significantly predicted individual MoCA scores. Critically, a model using only SUVR levels failed to predict MoCA scores. This demonstrates that dynamic brain measures capture a unique and clinically relevant signal of predicting cognitive impairments. The model’s top feature, increased S2 occupancy, predicted better cognition, directly linking the temporal properties of brain states to clinical function. This finding suggests that brain state dynamics may serve as an early and sensitive marker of the functional impact of emerging amyloid pathology, detecting breakdowns in neural efficiency before amyloid burden becomes sufficient to predict cognitive decline through static measures alone.

Collectively, our findings highlight the value of brain state dynamics for detecting subtle, functionally meaningful changes along the Aβ accumulation continuum. A key result is that while Aβ burden did not predict clinical diagnoses, symptoms, or task performance, variability in latent state dynamics successfully tracked both biological markers and cognitive outcomes. Latent state dynamics therefore offer a sensitive window into early disease processes. We conclude that this dimensional approach is a powerful tool for capturing continuous brain changes and individual differences that may precede overt cognitive impairment.

## Limitations and future directions

Several limitations warrant consideration. First, a key limitation is the investigation of Aβ pathology in a clinically heterogeneous sample. The absence of significant group differences in global Aβ SUVRs, particularly within our MCI cohort, suggests a notable portion of participants may have non-AD etiologies driving their cognitive deficits. While this complicates the interpretation of AD-specific pathology, our dimensional approach partially mitigates this by showing that brain dynamics relate to Aβ burden when it is present, and to cognition across the full sample regardless of etiology. Nonetheless, future studies must directly address this heterogeneity by stratifying analyses based on biomarker status (e.g., Aβ+ versus Aβ-MCI) to confirm the specificity of our findings to the Aβ continuum. Second, the analysis of brain state dynamics was restricted to a predefined set of network nodes, preventing a full whole-brain characterization of the latent states. This node-based approach was a necessary methodological constraint imposed by the substantial computational cost of modeling latent dynamics, which scales non-linearly with the number of regions. While our selected nodes were theoretically driven, capturing key large-scale networks involved in cognitive control, salience, default mode, and sensory networks, we acknowledge that regions outside this set may also critically contribute to the observed dynamics. Future work, enabled by advances in high-performance computing or more scalable algorithms, will be essential for extending these analyses to a whole-brain resolution. Last, the cross-sectional design precludes causal inferences and understanding of the longitudinal evolution of these dynamic changes. Future longitudinal studies are required to determine if these brain states can predict cognitive decline. Critically, such longitudinal work must also incorporate tau pathology, a core factor in AD with a distinct trajectory from Aβ (Callow et al., 2025; Chen et al., 2020; Chen et al., 2025; Gallego-Rudolf et al., 2024; Giacobini & Gold, 2013; Maass et al., 2018; Tideman et al., 2022). It remains an important open question whether and how these latent brain states are sensitive to tau burden. Multimodal longitudinal studies are therefore necessary to disentangle the unique contributions of Aβ and tau to these dynamic functional changes and their joint impact on cognitive impairment.

## Conclusion

In conclusion, this study demonstrates that latent brain state dynamics are a highly sensitive marker of early Aβ burden, succeeding where conventional metrics fail. In this cohort with predominantly subthreshold Aβ levels, amyloid burden itself did not predict clinical status, cognitive performance, or traditional static neural activation. In stark contrast, the temporal properties of latent brain states not only correlated with Aβ accumulation but also robustly predicted clinical MoCA scores and cognition. Critically, we identified a profound functional disconnection in MCI: while cognitively normal individuals showed a tight link between latent brain states engagement and task performance, this adaptive brain-behavior relationship was disrupted in the MCI group. This work establishes brain state dynamics as a powerful functional biomarker, offering a sensitive window into the neural impact of early pathology and providing a new avenue for identifying at-risk individuals.

## Methods

### Datasets

The Connectomics in Brain Aging and Dementia study (HCP-CBA, https://www.humanconnectome.org/study/connectomics-brain-aging-and-dementia) include 121 CN (74 F/37 M), 93 with impaired cognition (61 F/32 M) (Cohen et al., 2021). Each participant completed sessions of neuropsychological assessment, structural and functional neuroimaging and amyloid PET scans. During the functional neuroimaging session, participants underwent two runs of n-back working memory task. All participants provided written consent, and the local Institutional Review Board approved all study protocols.

### Data screen

Participants were selected based on their clinical diagnosis and data quality. Imaging data inclusion criteria was applied on both fMRI runs, including a mean framewise displacement (FD) of less than 0.3 mm per run and a maximum FD of less than 3 mm per run. This yielded an fMRI sample of 116 participants.

Behavioral data was available for 152 participants, comprising 78 cognitively normal participants and 74 MCI. Among the 116 participants whose neuroimaging data passes quality check, 14 lacked behavioral data for the n-back task, resulting in 102 participants for the combined fMRI and behavioral analysis. This analysis sample included 66 cognitively normal (CN) participants (44 F, 22 M; 63.47 years) and 36 individuals with mild cognitive impairment (MCI) (27 F, 9 M; 62.57 years).

### N-back task

The HCP n-back working memory task combines the category specific representation task and the n-back working memory task in a single task paradigm. Subjects were presented with blocks of trials that consisted of pictures of faces, places, tools and body parts. Within each session, the 4 different stimulus types were presented in separate blocks. Furthermore, within each session, half of the blocks are 2-back working memory and half are 0-back working memory task. In the 2-back working memory task blocks, subjects were requested to determine whether the current stimulus matches the stimulus in two presentations of stimuli prior within the same block. In the 0-back working memory task blocks, subjects were requested to determine whether the current stimulus matches the target that was presented in the beginning of each block (cue). A 2.5 second cue indicates the task type (and target for 0-back task) at the beginning of each block. Each of the two sessions contains 8 task blocks (10 trials of 2.5 seconds each, for 25 seconds) and 4 fixation (“rest”) blocks (15 seconds). On each trial, the stimulus is presented for 2 seconds, followed by a 0.5 second inter-trial-interval (ITI).

### Diagnostic Evaluation

Participants underwent a brief neuropsychological test battery designed for diagnostic classification, based on the Alzheimer’s Disease Research Center (ADRC) protocol. The battery included the Montreal Cognitive Assessment (MoCA) (Nasreddine et al., 2005) , which provides a global screening measure of cognitive functioning across multiple domains—attention, executive function, memory, language, visuospatial abilities, conceptual thinking, calculations, and orientation. Additional tests included verbal fluency, a 30-item confrontation naming test (Saxton et al., 2000), Trail Making Test (Reitan & Wolfson, 1994), verbal free recall (Welsh et al., 1991; Welsh et al., 1994), and the Rey–Osterrieth Complex Figure (Rey, 1941). Diagnostic classifications followed the ADRC scheme (Lopez et al., 2000). CN participants included those who reported no cognitive limitations as well as those with subjective cognitive complaints but without clinical diagnosis. The MCI includes two subgroups: individuals with impairment but no reported concerns (Impaired Without Complaints, IWC) and those who reported perceived loss of abilities, consistent with mild cognitive impairment (Cohen et al., 2013). For this study, we adopted the simplified division of participants into CN and MCI groups.

### MRI and Amyloid PET acquisition

Image protocol details can be found in a previous study (Cohen et al., 2021). The procedures are summarized here.

High-resolution structural and functional imaging was performed on a 3-Tesla Siemens Prisma scanner equipped with a 64-channel head coil. Anatomical images were acquired using a T1-weighted MP-RAGE sequence with an echo time (TE) of 2.22 ms, a flip angle of 8 degrees, and 0.8 mm isotropic voxels. Functional data for the N-back task were acquired using a high-temporal-resolution of 800 ms, gradient-echo EPI sequence with an echo time (TE) of 37 ms, flip angle of 52 degrees, 2.0 mm isotropic voxels.

Amyloid-β (Aβ) pathology was quantified using Positron Emission Tomography (PET) with the Pittsburgh Compound B (PiB) (Price et al., 2005; Wilson et al., 2000). Following a 20-second intravenous injection of approximately 15 mCi of high-specific-activity PiB (>0.50 Ci/µmol), participants rested for ∼25 minutes before being positioned in a Siemens/CTI ECAT HR+ scanner. A 30-minute PiB PET study (6 × 300 s frames) was acquired following a 10-minute transmission scan for attenuation correction. Data were reconstructed using filtered back-projection with a 3 mm FWHM Hann filter and corrected for scatter and radioactive decay, yielding a final image resolution of approximately 6 mm.

### fMRI preprocessing

A standard preprocessing pipeline was implemented using SPM12 software package (http://www.fil.ion.ucl.ac.uk/spm/software/spm12/), as well as in-house programs in MATLAB (MathWorks). Functional MRI data were first slice time corrected, aligned to the averaged time frame to correct for head motion, and co-registered with each participant’s T1-weighted images. Structural MRI images were segmented into grey matter, white matter, and cerebrospinal fluid. Based on the transformation matrix from structural image, the functional images were then transformed to the standard Montreal Neurological Institute (MNI) template in 2x2x2 mm^3^. A 6 mm Gaussian kernel was used to spatially smooth the functional images.

### Amyloid PET processing

Amyloid PET measures (regional and global) were obtained from preprocessed data publicly released by the University of Pittsburgh team. PET data were processed using PMOD and Freesurfer software. Following frame-to-frame motion correction, PET data were averaged to create a single image representing tracer uptake from 50-70 minutes post-injection. This image was then co-registered to the individual’s T1-weighted MRI. Cortical and subcortical regions of interest (ROIs) were automatically defined on the T1 image using Freesurfer’s segmentation and parcellation tools. Standardized uptake value ratios (SUVRs) were computed for the following regions of interest using a cerebellum reference region: anterior cingulate cortex, posterior cingulate cortex, superior frontal cortex, orbital frontal cortex, insular cortex, lateral temporal cortex, parietal lobe, precuneus, anterior ventral striatum, and averaged across hemispheres. The volume-weighted average of these SUVR values constituted the Global SUVR. More details can be found in a previous study (Cohen et al., 2021; Cohen et al., 2013).

### Regions of interest (ROI)

ROIs were determined by an independent study that investigates brain state dynamics in the Human Connectome Project (HCP) n-back working memory task (Taghia et al., 2018). The HCP and HCP-CBA studies have the same n-back working memory task. We selected 11 load-dependent ROIs by the contrast of interest: 2-back versus 0-back, including 9 load-positive (2-back > 0-back) ROIs, i.e. bilateral anterior insula (AI), bilateral middle frontal gyrus (MFG), bilateral frontal eye field (FEF), bilateral intraparietal sulcus (IPS) and dorsomedial prefrontal cortex (DMPFC), and 2 load-negative (2-back < 0-back) ROIs, i.e. ventromedial prefrontal cortex (VMPFC) and posterior cingulate cortex (PCC). In addition to these cognitive control network ROIs, eight category-specific visual ROIs (four for each hemisphere, corresponding to faces, places, tools, and body parts used in the task stimuli) were included to capture sensory processing aspects. The stimulus selectivity of these regions was confirmed using univariate contrasts and Neurosynth meta-analyses. Together, these 19 ROIs enable a comprehensive analysis of dynamic state transitions across salience, frontoparietal, default mode and visual networks (**Figure 2A**), with MNI coordinates provided in **Supplementary Table S1**.

### ROI time series

Each ROI was 6-mm radius sphere centered at the corresponding peak voxel. Mean activation of the time series was extracted from each ROI. A multiple linear regression approach with 6 realignment parameters (3 translations and 3 rotations) was applied to time series to reduce head-motion-related artifacts and resulting time series was further linearly detrended and normalized.

### Bayesian switching dynamical systems (BSDS) model

We applied BSDS, a generative state-space model, to identify latent brain states and their dynamic spatiotemporal signatures from the combined cohort time-series. In BSDS, the observed multivariate BOLD time-series are assumed to switch among a finite number of discrete latent states; each of which is characterized by its own low-dimensional continuous dynamical process and observation model. The switching among states is governed by a hidden Markov chain. The model allows the number of effective states to be estimated from the data under a variational Bayesian (VB) framework (Jordan et al., 1999; Wainwright & Jordan, 2008), avoiding reliance on arbitrary sliding-windows or predetermined task-locked intervals. In our implementation, time series data from all participants were combined and analyzed jointly to identify a common set of brain states across the entire cohort. The initial value for the number of possible latent brain states was set as 15. Details of the method can be found in the Supplementary Methods and our previous study (Cai et al., 2023; Cai et al., 2024; Taghia et al., 2018). Measures extracted from BSDS include occupancy rate and mean lifetime of latent brain state. The occupancy of each state is the amount of time spent in the state, which is computed by counting the number of time points in which a given state occurs. The mean lifetime of a state is the average time for which that particular state is continuously present and is computed by counting how long that state continuously persists and then taking an average of those counts.

### Multivariate Canonical Correlation Analysis (CCA)

We applied canonical correlation analysis (CCA) to assess multivariate relationships between regional SUVRs and behavioral as well as neural measures. First, we examined associations between regional SUVRs (X) and cognitive/task performance measures (Y), including accuracy, mean reaction time, and reaction time variability from both 0-back and 2-back conditions, as well as MoCA scores. Age and sex were included as nuisance covariates. Next, we tested the multivariate relationship between regional SUVRs and traditional univariate task activation. Activation estimates for the 0-back and 2-back conditions from 19 regions of interest (ROIs) defined in the BSDS analysis were entered as the Y variable set, with SUVRs as X. Age, sex, and mean head motion were included as additional covariates. Finally, we used CCA to relate regional SUVRs to latent brain state dynamics. Temporal features derived from the state analysis, such as task-dependent occupancy rate and mean lifetime, served as the X variable set, with regional SUVRs as Y; age, sex, and mean head motion were included as additional covariates. CCA and its statistical testing were conducted using ‘permcca’ algorithm implemented in Matlab. The ‘permcca’ algorithm was developed to address the concerns about shared variation on nuisance variables and repeatedly explained variations between canonical variables (Winkler et al., 2020). Permutation tests (1000 times) were used to build a null distribution of canonical correlations from which statistical significance were defined by *p*<0.05, corrected with the familywise error rate (FWER). The permutation test for CCA involves randomly shuffling the rows of either X or Y variables. With each permutation of the data denoted by π, a fresh set of canonical correlation r_π_ would be calculated. Subsequently, a p-value could be computed as *p*=(∑I(r_π_ >r_0_))/n_p_, where n_p_ is the number of permutations, r_0_ is the canonical correlation from non-permuted (original) data, and I = 1 if r_π_ >r_0_, otherwise I = 0. Last, *p* values were FWER-corrected for all the canonical correlation tests. Canonical coefficients were reported to characterize the multivariate relationship between two sets of variables. The contribution of each original variable on the canonical scores was assessed using Pearson’s correlation.

### Prediction models and cross validation

To test the robustness and generalizability of our findings, we built a series of predictive models using LASSO regression model. To ensure that our models could generalize to unseen data and to avoid overfitting, we employed a rigorous leave-one-out cross-validation (LOOCV) framework for all analyses. In this procedure, the model is iteratively trained on all participants except for one (the held-out participant) and then tested on its ability to predict the outcome for that left-out individual. This process is repeated until every participant has served as the test case once. The overall performance of the model was evaluated by calculating the Pearson’s correlation between the observed scores and the cross-validated predicted scores across all participants.

We investigated whether the temporal features of latent brain states could predict out-of-scanner clinical symptoms, and how their predictive power compared to a conventional biomarker of Aβ pathology. The target variable was the MoCA score. We trained and compared two separate models: a dynamics-only model: Using the mean lifetime and occupancy rates of the latent brain states as features; an Aβ-only model: Using the multi-regional Aβ SUVRs as features. This comparative approach allowed us to directly test whether dynamic brain measures provide unique, clinically relevant information beyond that captured by static amyloid PET measures.

## Acknowledgements

This research was supported by Alzheimer’s Association grants AARGD-NTF-21-850781 (WC), AARFD-21-849349 (CBY), AARFD-21-848178 (BL), and NIH grants EB022907(VM), NS086085 (VM), and R00AG071837 (CBY), and the Wu Tsai Neurosciences Institute at Stanford University (VM).

## Author contributions

Study design: W.C., V.M.; Data Analysis: Z.G., C.B.Y., B.L., W.C.; Manuscript Drafting: Z.G., W.C.; Manuscript Editing: Z.G., C.B.Y, B.L., V.M., W.C.

## Competing financial interests

C.B.Y. has served as a consultant to Medidata Solutions. E.C.M. has served as a consultant for Roche, Genentech, Eli Lilly, and Neurotrack. All other authors declare no competing financial interests.

