## Supplementary Materials for "Latent brain state dynamics predict early amyloid accumulation and cognitive impairment"

**Method**

**Bayesian Switching Dynamical Systems (BSDS) Generative Model**

To investigate latent brain state dynamics, we applied a Bayesian Switching Dynamical Systems (BSDS) model, a state-space generative framework designed to uncover shared brain states across multiple cognitive tasks. This model identifies brain states and their dynamic spatiotemporal properties within an optimal latent subspace. Let $\boldsymbol{y}_{t}^{s}$ denote a $D$-dimensional vector of observed fMRI measurements in time $t$ and for subject $s$, and $\boldsymbol{z}_{t}^{s}$ denote a $1$-of-$K$discrete vector of latent state variables of a hidden Markov model (HMM) with elements $z_{kt}^{s}, \forall k=1,\ldots,K.$Two consecutive time instances are dependent via a first-order Markov chain through an HMM. Specifically, the probability distribution of $\boldsymbol{z}_{t}^{s}$ depends on previous latent variable $\boldsymbol{z}_{t-1}^{s}$through a conditional distribution $p\left( \boldsymbol{z}_{t}^{s} \mid\boldsymbol{z}_{t-1}^{s}\boldsymbol{,A} \right)= \prod_{k=1}^{K} {\prod_{j=1}^{K} A}_{jk}^{z_{t-1,j}^{s} z_{tk}^{s}}$ for all $t>1$ represented by the transition probabilities $\boldsymbol{A}$**,** where $A_{jk}\equiv p\left( z_{tk}^{s}=1 \mid z_{t-1,j}^{s}=1 \right)$, and a marginal distribution $p\left( \boldsymbol{z}_{1}^{s}\mid\boldsymbol{\pi} \right)=\prod_{k=1}^{K} \pi_{k}^{z_{1k}^{s}}$ represented by a vector of initial probabilities $\boldsymbol{\pi}$ where $\pi_{k}\equiv p\left( z_{1k}^{s}=1 \right)$ (Bishop & Nasrabadi, 2006). Next, we assume that at a given mode of the system given by the latent state $z_{kt}^{s}=1$, observed vector $\boldsymbol{y}_{t}^{s}$ is generated via a state space model in form of$\mathbf{:}$

$\boldsymbol{y}_{t}^{s}=\boldsymbol{U}_{k}\boldsymbol{x}_{kt}^{s}+\boldsymbol{\mu}_{k}\boldsymbol{+}e_{kt}, \forall t\mid z_{kt}^{s}=1,$	$\boldsymbol{x}_{kt}^{s}={\bar{\boldsymbol{X}}}_{kt}^{\boldsymbol{s}} {\vec{\boldsymbol{V}}}_{k}\boldsymbol{+}\boldsymbol{\epsilon}_{kt}\boldsymbol{,}\boldsymbol{\forall}t\mid z_{kt}^{s}=1.$	(1)

The first line of the generative model in Eq. (1) can be viewed as a probabilistic factor analysis model where $\boldsymbol{U}_{k}$ is a $D\times P$dimensional linear transformation matrix that transforms data to a subspace of lower dimensionality, $P<D$, described using a $P$-dimensional vector of latent space variables $\boldsymbol{x}_{kt}^{s}$ mediated by an overall bias $\boldsymbol{\mu}_{k}$ and a measurement noise $\boldsymbol{e}_{kt}\boldsymbol{\sim}\mathcal{N(}\boldsymbol{0,}\boldsymbol{\Psi}_{\boldsymbol{k}})$ (Everett, 2013; Ghahramani & Beal, 1999). The latent dynamics of the system are further modeled with an autoregressive (AR) process of order $R$ defined on the latent space variables of the factor analysis model (Fox et al., 2008). ${\vec{\boldsymbol{V}}}_{k}$ is a vector of AR coefficients$.$ ${\bar{\boldsymbol{X}}}_{kt}^{\boldsymbol{s}}\boldsymbol{=}{\mathrm{diag}(\bar{\boldsymbol{x}}}_{kt}^{\boldsymbol{s}}\boldsymbol{)}$ is a block diagonal isotropic matrix with elements of ${\bar{\boldsymbol{x}}}_{kt}^{\boldsymbol{s}}\boldsymbol{=(}{\boldsymbol{x}_{k,t-1}^{s}}^{T}\boldsymbol{,}{\boldsymbol{x}_{k,t-2}^{s}}^{T}\boldsymbol{,}\ldots, {\boldsymbol{x}_{k,t-R}^{s}}^{T}\boldsymbol{)}$ represented using latent space variables from the previous $R$ time frames where T indicates the transpose operator. $\boldsymbol{\epsilon}_{kt}\mathcal{\sim N(}\boldsymbol{m}_{k}, \boldsymbol{\Sigma}_{k})$ models the remaining error term in the latent space. An AR process of a first order, $R=1,$ is defined on the representations of the observations in the latent subspace, $\boldsymbol{x}_{kt}^{s}$. This method provides a powerful framework for identifying spatiotemporal patterns associated with latent brain states. Detailed theoretical derivations for this model are available in previous studies (Taghia et al., 2018).

**Supplementary Results**

**Brain states are differently sensitive to WM loads**

To understand how latent brain state dynamics relate to task demands and clinical status, we conducted group-level analyses. Specifically, we tested whether mean lifetime and occupancy rate of brain states were modulated by working memory load and whether there is interaction effect between working memory load and clinical groups.

We conducted an ANOVA with two within-subject factors, i.e. load (low and high) and state (S1, S2, S3 and S4), and one between-subject factor (MCI and CN). The ANOVA on mean lifetime revealed no significant three-way interaction (F(1.929, 167.04) = 0.217, p = 0.765, η^2^_p_ = 0.002, Greenhouse-Geisser corrected). However, there was a significant interaction between WM Load and State (F(1.167, 167.04) = 53.596, p < 0.001, η^2^_p_ = 0.349, Greenhouse-Geisser corrected), suggesting WM load significantly modulated brain state dynamics. There was also a significant main effect of State (2.012,201.199) = 459.760, η^2^_p_ = 0.821, Greenhouse-Geisser corrected), but the main effect of WM Load was not significant (F(1, 100) = 0.133, p = 0.716, η^2^_p_ = 0.001) (Figure 3A). Critically, there was no significant main effect of Group (F(1,100) = 0.017, p = 0.897, η^2^_p_ < 0.001). Furthermore, none of the interactions involving the Group factor were significant: WM Load × Group (F(1, 100) = 0.350, p = 0.555, η^2^_p_ = .003), Brain State × Group (F(2.012, 201.199) = 0.474, p = 0.624, η^2^_p_ = 0.005, Greenhouse-Geisser corrected), and WM Load × Brain State × Group (F(1.670, 167.040) = 0.217, p = 0.765, η^2^_p_ = .002, Greenhouse-Geisser corrected). Similarly, the ANOVA for occupancy rate showed no significant three-way interaction (F(1.452, 145.237) = 0.557, p = 0.519, η^2^_p_ = 0.006, Greenhouse-Geisser corrected). There were significant main effects of State (F(1.738, 173.753) = 354.988, p<.001, η^2^_p_ = 0.780, Greenhouse-Geisser corrected) and a significant WM Load × State interaction (F(1.452, 145.237) = 86.001, p < 0.001, η^2^_p_ = 0.462), but no significant main effect of WM Load (F(1,100) = 2.042, p=0.156, η2p < 0.020). Consistent with the mean lifetime results, there was no significant main effect of Group (F(1,100) = 0.015, p = 0.904, η^2^_p_ < 0.001), nor were any interactions involving Group significant: WM Load × Group (F(1,100) = 3.837, p = 0.053, η^2^_p_ = 0.037), State × Group (F(1.738, 173.753) = 1.200, p = 0.299, η^2^_p_ = 0.012, Greenhouse-Geisser corrected), and WM Load × State × Group (F(1.452, 145.237) = 0.557, p = 0.519, η^2^_p_ = 0.006, Greenhouse-Geisser corrected).

Given that no significant interactions involving groups were detected for either mean lifetime or occupancy rate, subsequent analyses were conducted by collapsing across groups to examine the effects of working memory load on each brain state. Simple effects analysis revealed that states S2 and S3 exhibited significantly higher mean lifetime and occupancy rate during 0-back trials compared to 2-back trials (all ps < 0.001, FDR-corrected for each measure separately). The state S2 had significantly higher mean lifetime and occupancy rate than all the other states during 0-back trials (all ps < 0.001, FDR-corrected). This suggests that the state S2 is the dominated state in low-load (0-back) condition. In contrast, state S4 showed the opposite pattern, with significantly higher mean lifetime and occupancy rate during 2-back trials relative to 0-back trials (all ps < 0.001, FDR-corrected for each measure separately). The state S4 had significantly higher mean lifetime and occupancy rate than all the other states during 2-back trials (occupancy rate: S4 vs. S2, p = 0.054; all other ps < 0.001, FDR-corrected). This suggests that the state S4 is the dominated state in high-load (2-back) condition. Meanwhile, state S1 did not show significant modulation by working memory load on either measure (mean lifetime: p = 0.390; occupancy rate: p = 0.202; FDR-corrected). Overall, these results indicate that specific brain states are differentially sensitive to working memory demands, implying distinct functional roles across cognitive load conditions.

**Brain state dynamics predict task performance using a dimensional approach**

We also examined brain-behavior relationships across all participants using a dimensional approach, controlling for age, sex, and head motion. Under the low‐load (0‐back) condition, distinct brain–behavior relationships emerged. Longer mean lifetimes of states S2 and S3 were associated with higher efficiency (r = 0.242, 0.224; p = 0.033, 0.038) and accuracy (r = 0.270, 0.275; p = 0.011, 0.011), as well as faster responses (r = −0.205, −0.226; p = 0.057, 0.054). Neither state was significantly related to response‐time variability (r = −0.135, −0.184; p = 0.245, 0.150). In contrast, longer S4 lifetimes predicted lower efficiency (r = −0.416; p < 0.001) and accuracy (r = −0.466; p < 0.001), slower responses (r = 0.358; p = 0.001), and greater response‐time variability (r = 0.268; p = 0.033). Occupancy rates showed a parallel pattern: greater S2 and S3 occupancy was linked to higher efficiency (r = 0.281, 0.237; p = 0.010, 0.028) and accuracy (r = 0.323, 0.258; p = 0.003, 0.017), and faster responses (r = −0.217, −0.259; p = 0.043, 0.023), S3 also negatively associated with response-time variability (r = -0.233, p = 0.048); higher S4 occupancy predicted lower efficiency (r = −0.459; p < 0.001), lower accuracy (r = −0.504; p < 0.001), slower responses (r = 0.393; p < 0.001), and greater response‐time variability (r = 0.269; p = 0.030). All p values were FDR‐corrected for four brain states. Under the high‐load (2‐back) condition, neither lifetime nor occupancy rates significantly predicted efficiency, accuracy, speed, or variability (all p > 0.133, FDR corrected).

**Distinct brain activation and connectivity patterns underlying latent brain states**

Our analyses reveal that latent brain states not only strongly predict amyloid-beta levels but also play a critical role in cognitive function. In particular, states S2 and S4 were the dominated states in the low-load and high-load conditions, respectively. To understand the underlying neural mechanisms, we investigated the distinctive patterns of regional activation and functional connectivity (FC) that define these states.

A direct comparison between the two states revealed robust activation differences across nearly all ROIs. Relative to the State S2, the State S4 showed significantly greater activation in salience and frontoparietal network regions, as well as in visual areas selective for place, the bilateral PHA (all ps < 0.001, FDR-corrected). In contrast, S4 exhibited stronger deactivation in default mode network regions and lower activation in visual areas selective for living categories such as faces and bodies, as well as the tool-selective region in the right MTG (all ps < 0.001, FDR-corrected); only the left MTG tool-selective region did not show a significant difference (p = 0.733), see Figure S1. For connectivity, relative to the State S2, the State S4 exhibited widespread hyperconnectivity, most prominently within and between the FPN, DMN, and SN (Figure S2). Together, these activation and connectivity patterns suggest that S4 reflects a high-load state marked by enhanced recruitment of cognitive control systems to support demanding task conditions.

Supplementary Figures

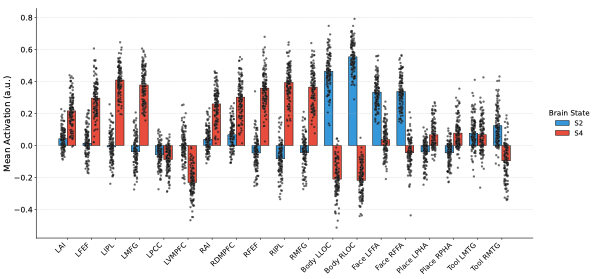

Figure S1. Brain regional activation differences between S2 and S4. Bilateral middle frontal gyrus (MFG), frontal eye field (FEF), inferior parietal lobe (IPL); bilateral anterior insular, dorsal medial frontal cortex (dmPFC); ventral medial frontal cortex (vmPFC) and posterior cingulate cortex (PCC), face fusiform area (FFA), place parahippocampus area (PPA), extrastriate body area (EBA), and tool selective area in MTG

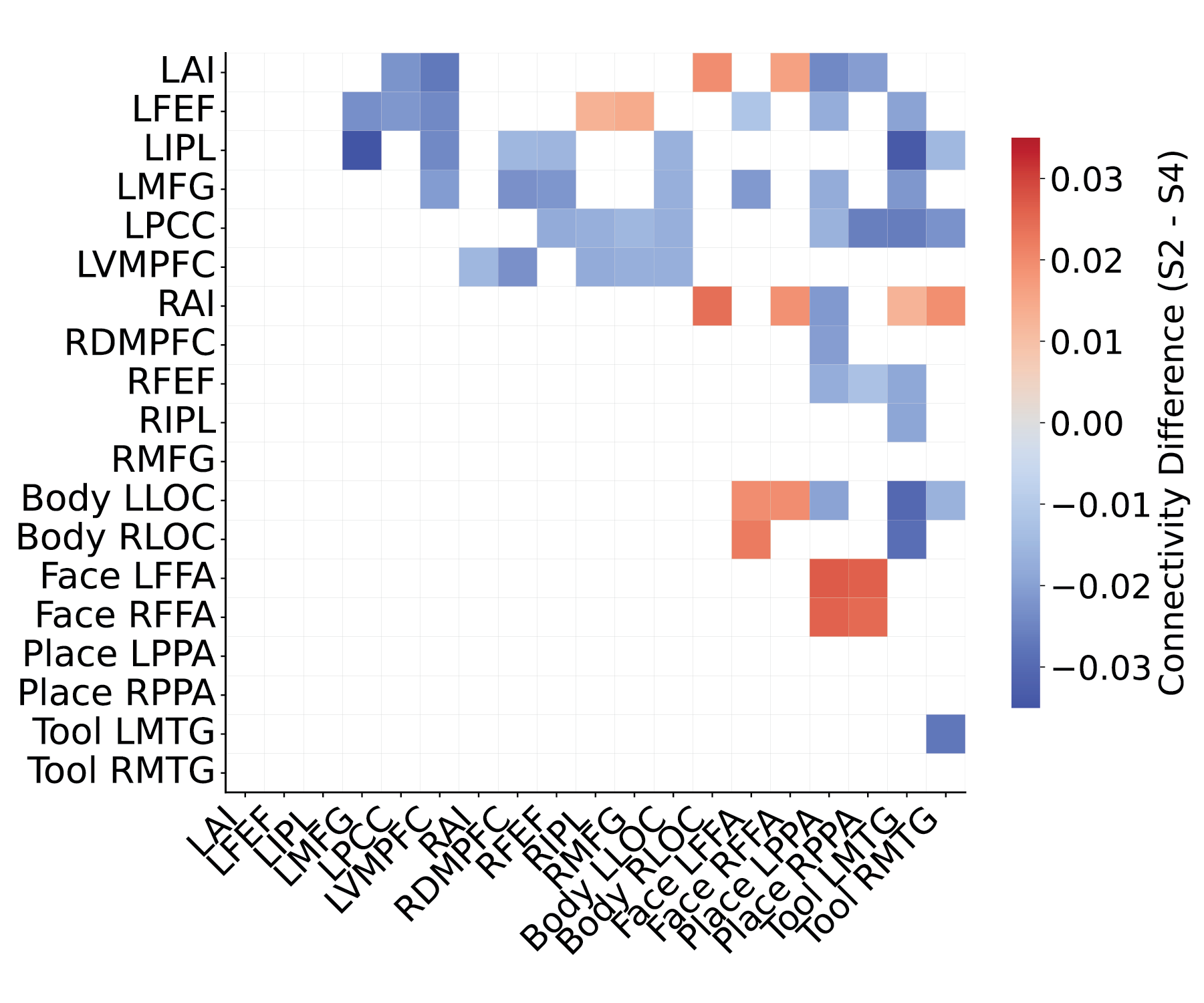

Figure S2. Brain connectivity differences between S2 and S4. Bilateral middle frontal gyrus (MFG), frontal eye field (FEF), inferior parietal lobe (IPL); bilateral anterior insular, dorsal medial frontal cortex (dmPFC); ventral medial frontal cortex (vmPFC) and posterior cingulate cortex (PCC), face fusiform area (FFA), place parahippocampus area (PPA), extrastriate body area (EBA), and tool selective area in MTG

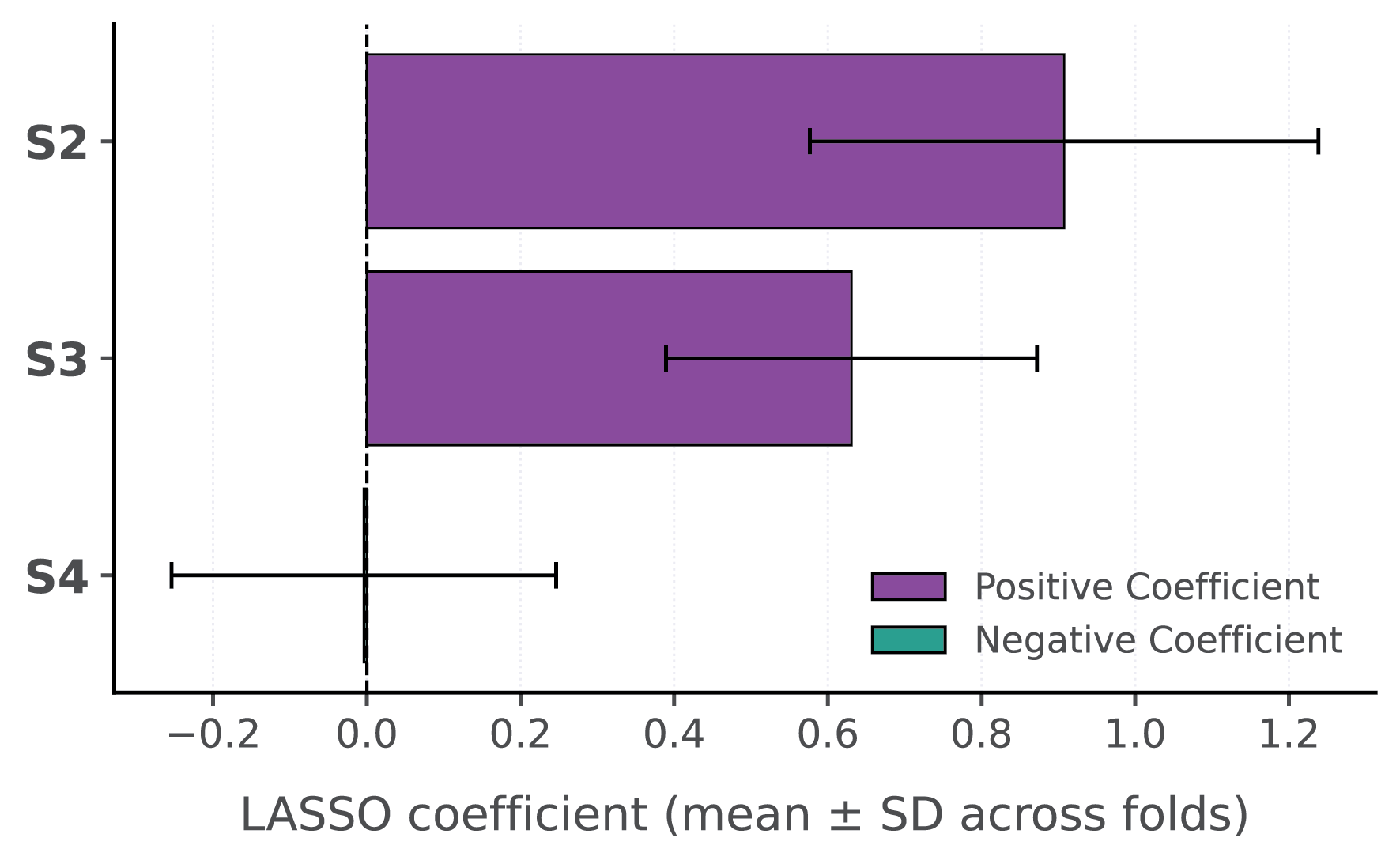

Figure S3. LASSO feature weights for MoCA score prediction using brain state occupancy. Shown are the mean coefficients (± SD across cross-validation folds) for a model predicting MoCA scores. Features were the occupancy rates of states S2, S3, and S4 (with S1 excluded to prevent multicollinearity). Positive coefficients are associated with higher MoCA scores.

| ROI | MNI Coordinate | | |
| --- | --- | --- | --- |
|  | X | Y | Z |
| LAI | -32 | 24 | 2 |
| LFEF | -26 | 2 | 58 |
| LIPS | -46 | -44 | 44 |
| LMFG | -42 | 24 | 30 |
| LPCC | -12 | -56 | 16 |
| LVMPFC | -2 | 48 | -8 |
| RAI | 36 | 22 | 0 |
| RDMPFC | 4 | 16 | 50 |
| RFEF | 30 | 10 | 56 |
| RIPS | 52 | -40 | 50 |
| RMFG | 40 | 36 | 34 |
| Body LLOC | -48 | -70 | 4 |
| Body RLOC | 48 | -68 | 0 |
| Face LFG | -40 | -54 | -20 |
| Face RFG | 44 | -48 | -20 |
| Place LLG | -26 | -48 | -10 |
| Place RLG | 26 | -44 | -10 |
| Tool LMTG | -54 | -54 | -6 |
| Tool RMTG | 54 | -54 | 10 |

Table S1. MNI coordinates of the 19 ROIs used for dynamic state transition analysis. Coordinates are provided for 11 load-dependent ROIs (bilateral AI, MFG, FEF, IPS, DMPFC, VMPFC, and PCC) selected via 2-back versus 0-back contrasts (Taghia et al., 2018), and 8 stimulus-selective visual ROIs. Together, these regions capture activity across salience, frontoparietal, default mode, and visual networks.
